# Central and peripheral dynamics of acute stress: evidence from functional cortical gradients

**DOI:** 10.64898/2026.07.08.732866

**Authors:** Agata Patyczek, Elias Reinwarth, Janis Reinelt, Arno Villringer, Marie Uhlig, Samyogita Hardikar, Michael Gaebler

## Abstract

Stress involves coordinated central and peripheral processes that unfold dynamically and can be assessed through brain, autonomic, endocrine, and subjective measures. Centrally, acute stress has been linked to altered functional connectivity, particularly in the salience (SN), frontoparietal networks (FPN), and default mode networks (DMN). Here, we used cortical gradients to characterize stress-related reconfiguration in macroscale functional space and assessed their relation to peripheral stress measures.

We performed secondary analyses on data from 67 young males completing the Trier Social Stress Test or a control task with resting-state fMRI before and after, concurrent peripheral (autonomic, endocrine) and subjective measures. To assess region- and network-specific changes in functional organization, we derived eccentricity and within- and between-network dispersion for the first three cortical gradients.

Acute stress was associated with selective gradient reconfigurations in the right ventral prefrontal cortex and left insula and with increased SN–DMN and SN–FPN dispersion, indicating DMN and FPN decoupling from the SN. Although no associations with peripheral or subjective stress measures survived multiple-comparison correction, nominal effects suggested partly distinct links of saliva cortisol with local gradient changes and HRV with network-level reconfiguration. Together, these findings show that acute stress selectively reconfigures macroscale cortical organization.

## Introduction

Short-term or acute stress activates a set of physiological and psychological changes to adaptively respond to the stressor (1). While the acute stress response is adaptive, prolonged or chronic stress increases the risk of physical and mental health disorders (2).

Acute stress can alter brain activity in regions such as the amygdala, prefrontal cortex, hippocampus, and insula, which are also related to emotional regulation, cognitive control, and memory (3–5). Extending to connectivity, the “tri-network” hypothesis (6) proposes that stress drives a dynamic reallocation of resources among three key networks: the salience network (SN), the frontoparietal (FPN; sometimes referred to as the central executive network), and the default mode network (DMN). Psychologically, heightened SN activity has been associated with greater vigilance (6,7), decreased DMN activity to reduced self-referential thought (8,9), and DMN–FPN decoupling to diminished cognitive control (10,11).

Adaptations to acute stress extend beyond the brain to other bodily systems, most prominently the autonomic nervous system (ANS), which adjusts cardiovascular function through rapid parasympathetic withdrawal and sympathetic activation, giving rise to measurable changes in heart rate and heart rate variability, and the hypothalamic-pituitary-adrenal (HPA) axis, which initiates a slower hormonal cascade in which adrenocorticotropic hormone (ACTH) release precedes cortisol secretion, which can be measured peripherally in plasma (e.g., ACTH) or saliva (e.g., cortisol) (12). Acute stress is also subjectively experienced, shaping how individuals appraise and react to a stressor. We previously showed that peripheral and subjective stress measures are differentially related to stress-induced brain changes. For example, salivary cortisone correlated with state anxiety and heart rate (13); stress-related thalamic functional connectivity was associated with subjective stress and, more weakly, with HRV and salivary cortisol (14); and stress-related gray matter volume changes were significantly related to HRV and state anxiety, but not to endocrine measures (15). Together, these findings support that acute stress constitutes an overarching state of organism-wide adaptation, in which peripheral, subjective, and brain responses co-vary, but capture partly distinct facets of the stress response.

To capture distributed stress-related changes in the brain and relate them to peripheral and subjective stress measures is supported by analysis approaches that simultaneously consider macroscale characteristics (beyond discrete areas or parcellations). We here used cortical gradient analysis, which is particularly well suited to study acute stress in the brain and relate it to other stress measures.

Gradient approaches characterize cortical organization along continuous, low-dimensional axes of differentiation that reflect systematic transitions in cytoarchitectural complexity, laminar organization, and functional specialization across the cortical sheet (16–18). The principal cortical gradient captures, for example, how cortical regions range from highly granular, myelinated sensory and motor areas with sharply defined laminar structure to less architecturally differentiated, transmodal association regions - a continuum that reflects the brain’s functional spectrum from stimulus-bound processing to abstract, integrative cognition (16). Additional gradients differentiate sensory modalities (e.g., visual from somatomotor cortex) and resolve further distinctions among transmodal systems, including the SN, FPN, and DMN (17,18).

In this study, we examined how acute psychosocial stress alters cortical functional organization by mapping stress-related connectivity changes onto the brain’s macroscale functional axes using cortical gradients. We focused on the cortex because cortical gradients have been most extensively characterized and reliably replicated. To assess stress-related changes in cortical organization, we preregistered (https://osf.io/5yspn) and performed a cortex-wide analysis in gradient space, using eccentricity - reflecting the functional distinctiveness of individual brain regions within the cortical hierarchy - and network dispersion - capturing the degree of functional differentiation within and segregation between large-scale networks. This approach allowed us to investigate both regional connectivity shifts (eccentricity) and network-level reconfiguration (network dispersion), with a particular focus on the three networks implied in the stress-related reallocation of resources (6) - SN, FPN, and DMN. By examining both within-network and between-network changes, we aimed to capture stress-related alterations across the tri-network architecture more comprehensively. We then explored how stress-related gradient changes relate to peripheral and subjective stress measures, testing whether different facets of cortical reorganization map selectively onto autonomic, endocrine, and subjective experiential indices of stress reactivity.

## Methods

The following methods for the secondary data analysis were preregistered and can be found at: https://osf.io/5yspn. Any analysis outside the initial preregistration is labelled as exploratory. As the focus of this paper is on dynamics, we here omit the preregistered analysis about chronic stress-related cortical gradients (to be published elsewhere).

### Participants

Data from the “Neural Consequences of Stress” dataset (NECOS: (13,14)) was used. NECOS is an extension of the LEMON study (19), where participants underwent an additional protocol to examine acute psychosocial stress. The study comprised 67 young male participants (aged 18-35) who were randomly assigned to either a stressor group (n=33) or a placebo/control group (n=34). To minimize the influence of diurnal variations on cortisol levels, data were collected during the same time of day for all participants, between 12:15 and 17:35. All participants provided written informed consent before their involvement in the study. Prior results and experimental details of NECOS can be found in prior publications (13–15). The study was approved by the ethics committee at the Medical Faculty of the University of Leipzig (number 385-1417112014).

### Procedure

Participants first underwent baseline preparation, including placement of an intravenous catheter and a portable ECG device, followed by repeated saliva, blood, and subjective assessments. After a standardised lunch, completion of trait questionnaires, and a short rest period, they completed the pre-intervention MRI session, which comprised a structural scan and two resting-state fMRI runs (rest1 and rest2), with additional in-scanner sampling between sequences. Importantly, the experimenters remained unaware of group assignment until after completion of rest2.

Acute psychosocial stress was induced using the Trier Social Stress Test (20). Participants faced a committee of two professional actors, introduced as trained psychologists skilled in nonverbal communication analysis. They were asked to present their qualifications for a desired job while being audio- and video-recorded. After a 5-minute preparation phase, they delivered the presentation without notes, while the committee remained neutral and interrupted only to repeat instructions if needed. Participants then performed a backward-counting task from 2043 in steps of 17. This procedure reliably induces stress, as indexed by saliva cortisol levels (13–15). To prolong and intensify the stress response, participants were told they would complete another task inside the MRI scanner and were then accompanied to the MRI area.

After the intervention, participants completed a post-intervention MRI session with four additional resting-state runs (rest3–rest6), a second structural scan, and repeated saliva, blood, and subjective assessments collected between sequences. After the second structural scan, stress-group participants were told that no further task would follow, marking the start of recovery. The session ended with a final assessment and debriefing. As a between-subject control condition, a placebo-TSST matched the TSST in timing, posture changes, speaking, and mental arithmetic, but omitted its socially evaluative and stress-inducing elements, such as critical observation by a committee. (21).

### Measures

#### Central stress measures

##### MRI acquisition and preprocessing

Magnetic resonance imaging (MRI) data was collected at the Max Planck Institute for Human Cognitive and Brain Sciences in Leipzig, Germany, using a 3T MAGNETOM Verio scanner (Siemens, Erlangen, Germany) with a 32-channel head coil. High-resolution structural images were obtained through a Magnetization Prepared 2 Rapid Acquisition Gradient Echoes (MP2RAGE) sequence, featuring a sagittal acquisition orientation, 176 slices, including a repetition time (TR) of 5000 ms, echo time (TE) of 2.92 ms, and inversion times (TI1/TI2) of 700 ms and 2500 ms, respectively. The flip angles were set at 4° and 5°, with a field of view (FOV) of 256 mm and a voxel size of 1 mm isotropic. Additional parameters included echo spacing of 6.9 ms, bandwidth of 240 Hz/pixel, GRAPPA acceleration factor of 3, and interleaved slice order.

Resting-state functional MRI (rs-fMRI) data were acquired using a T2*-weighted echo-planar imaging (EPI) sequence with an axial acquisition orientation. The rs-fMRI parameters included a voxel size of 2.3 mm isotropic, FOV of 202 mm, imaging matrix of 88 x 88, and 64 slices with a thickness of 2.3 mm. The TR was set to 1400 ms, TE to 30 ms, and the flip angle to 69°. Each rs-fMRI scan lasted 8.22 minutes. Before the rs-fMRI scans, field mapping was performed using gradient echo non-EPI scans and spin-echo EPI scans for distortion correction. Data preprocessing for the rs-fMRI was conducted using the FMRIB Software Library (FSL), while spatial transformations and structural preprocessing were carried out with Advanced Normalization Tools (ANTs). The complete pipeline was implemented using Nipype and can be found here: (https://github.com/NeuroanatomyAndConnectivity/pipelines/tree/master/src/lsd_lemon).

##### Gradient analysis

Functional cortical gradients during rest were computed using the BrainSpace toolbox (Version 0.1.4) in Python 3.9 (22) *(Figure 1A).* First, for each participant, a functional time series was extracted with a Schaefer 400 parcellation (23). From the resulting time series, a 400 by 400 connectivity matrix was calculated using Pearson correlation for every participant. Next, group connectivity matrices were created as the average of all participants. The affinity matrices were built using the normalized angle kernel (sparsity = 0.9, in line with (24)). The affinity matrices were then linearly decomposed using Principal Component Analysis (PCA), due to better reliability, compared to nonlinear methods, when predicting phenotypic scores (25).

**Figure 1.**
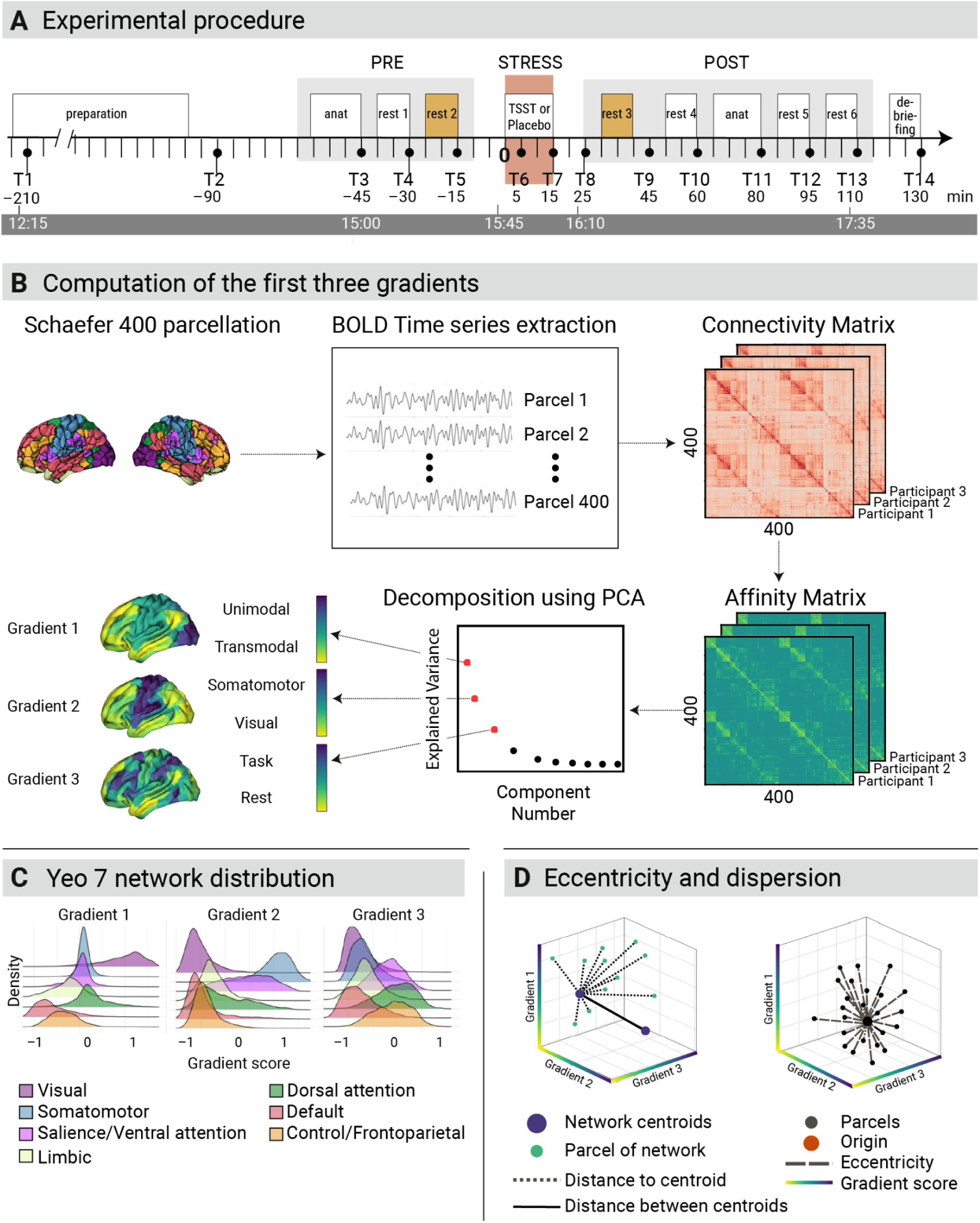
Experimental procedure, gradient derivation, network distribution, and spatial dispersion analyses. **(A)** Experimental procedure of NECOS. Over the course of the experiment (12:15-17:35 for all participants), six 8-min resting-state fMRI segments were acquired with concurrent Electrocardiography (ECG) and photoplethysmography (PPG) were recorded throughout the session, and ECG was additionally recorded outside the scanner during the Trier Social Stress Test (TSST) at three time points (not shown here). Two scans were acquired before the intervention (rest1 and rest2), whereas four scans were acquired after the intervention (rest3-rest6). The present analyses focused on the resting-state blocks immediately preceding (rest2) and following (rest3) the intervention (highlighted in orange). Blood samples were collected at 14 time points (T1-T14), and saliva samples and subjective ratings were obtained at 15 time points (T0-T14). **(B)** Workflow for computing the first three cortical connectivity gradients. BOLD time series were extracted from the Schaefer 400-parcel atlas, used to construct parcelwise connectivity matrices, transformed into affinity matrices, and decomposed using principal component analysis (PCA). The first three gradients are shown on the cortical surface and span canonical axes of functional organization, including unimodal-to-transmodal and sensory-to-task-related dimensions. (**C)** Distribution of gradient scores across Yeo’s seven functional networks for each of the first three gradients, illustrating how large-scale networks are positioned along each gradient axis. **(D)** Schematic of eccentricity and dispersion measures in three-dimensional gradient space. Network centroids were defined as the mean position of parcels belonging to each network; eccentricity reflects the distance of parcels from the origin, whereas dispersion reflects the spread of parcels relative to their network centroid.

To improve comparison with prior studies, the group gradients were aligned to the Human Connectome Project (HCP) dataset using the Procrustes alignment as provided by BrainSpace (24). The subsequent analysis resulted in a 400-by-3 (parcel-by-gradient) tensor. Finally, to attribute each parcel to a network, the Yeo 7 network atlas was used to assign the 400 parcels to the 7 canonical functional networks (26) *(Figure 1B).* A voxel-based exploratory comparison was conducted to identify the subdivisions within significant regions using the Harvard-Oxford and Brainnetome atlases (27,28).

##### Eccentricity and dispersion measure

Eccentricity measures how far each parcel lies from the center of the 3D gradient space, defined by the first three zero-centered gradients, one on each axis (29) (Figure 1C, right side). Each parcel’s position is determined by its scores on these gradients, and eccentricity is calculated as the Euclidean distance to the origin: the square root of the summed squared gradient values across the three axes. Higher eccentricity therefore indicates parcels farther from the center of the manifold space.

Dispersion measures how widely parcels are distributed in gradient space (Figure 1C, left). For each participant, we first calculated each network’s centroid as the mean position of its parcels in 3D gradient space. Within-network dispersion was defined as the sum of squared Euclidean distances between each parcel and its network centroid, producing one value per network per participant. Between-network dispersion was calculated as the Euclidean distance between pairs of network centroids. Both eccentricity and dispersion were computed for all six available resting states.

##### Peripheral and subjective stress measures

Peripheral and subjective stress measures used in this secondary analysis were collected repeatedly throughout the whole protocol in the same cohort, using a dense sampling scheme designed to capture autonomic, endocrine, and subjective stress dynamics before, during, and after stress exposure (13–15). Blood was sampled at 14 time points (T1–T14) via an indwelling cubital venous catheter, whereas saliva and subjective ratings were obtained at 15 time points (T0–T14); saliva was collected with Salivette devices (Sarstedt AG & Co. KG, Nümbrecht, Germany) for at least 2 min per sample, and blood and saliva samples were centrifuged and stored at −80 °C until analysis.

Subjective stress measures comprised repeated assessments of state anxiety using the state version of the State-Trait Anxiety Inventory (STAI; sum score of the state subscale) and perceived stress using the question “How stressed do you feel right now?”, rated on a visual analogue scale (VAS) from 0 (“not at all”) to 100 (“very much”). Questionnaires were presented both outside and inside the scanner using the same task framework across the sessions. Autonomic stress measures were recorded continuously throughout the experiment. Outside the scanner, heart activity was acquired with a BioHarness3 (Zephyr, Annapolis, Maryland, US) chest strap ECG; inside the scanner, ECG was recorded with BrainAmp, and pulse oximetry/photoplethysmography was recorded at the finger with an MP150 system. Heart rate and RMSSD-based heart rate variability were then derived in 3-min bins from interbeat intervals, with RMSSD indexing parasympathetic cardioregulation. Endocrine stress measures were quantified from the blood and saliva samples using LC-MS/MS-based assays as described in the original cohort reports (13–15).

### Statistical analysis

Statistical testing was conducted in R version 4.5.0 (30). Linear mixed-effects models (LMMs; lme4 package) tested effects of scan (rest2, rest3), group (stress, control), and their interaction on eccentricity and dispersion, using participant-specific random intercepts and likelihood-ratio tests for model comparison (31). Confirmatory post-hoc analyses of the three individual gradients were Bonferroni-corrected. To test whether individual physiological and subjective stress responses predicted pre-to-post changes in eccentricity and dispersion, we used delta ANCOVAs adjusted for baseline outcome values at rest2. For each brain outcome, change from rest2 to rest3 was modeled as a function of the group, the respective stress-reactivity measure, their interaction, the baseline brain outcome, and head motion (framewise displacement). These exploratory analyses were FDR-corrected. Within the stress group, Spearman correlations assessed associations between changes in peripheral stress measures and changes in eccentricity or dispersion, with FDR correction applied separately within each brain-measure family. Additional three-way random-intercept LMMs were estimated as uncorrected sensitivity analyses. All analysis scripts are available in the Git repository: (https://github.com/elrei-science/NECOS_gradients)

## Results

### Cortical gradients

The first three functional cortical gradients explained 67% of variance within the affinity matrix in pre (session 2) and 68% in post (session 3), similarly to previous studies (Margulies et al., 2016; Vos de Wael et al., 2020b). The principal gradient distinguished unimodal from transmodal regions, with lower values indicating higher similarity to visual regions and higher values indicating higher similarity to the DMN. The second gradient separated unimodal areas spanning from somatomotor to visual regions, while the third gradient delineated transmodal systems, contrasting the DMN to the FPN.

### Eccentricity

A significant interaction between scan session (pre-stress (rest2) vs. post-stress (rest3)) and group (stress vs. control) was identified in two key brain regions: the right ventral prefrontal cortex (Schaefer 400 ROI: 377, MNI centroid coordinates: 23, 55, 7; T = -4.84, pFDR = 0.003) and the left insula (Saefer 400 ROI: 100; MNI coordinates: -43, 4, -1; T = -3.90, pFDR = 0.045) (Figure 2A). Complete cross-atlas comparison results are reported in the Supplementary Material (Tables 1 and 2).

**Figure 2:**
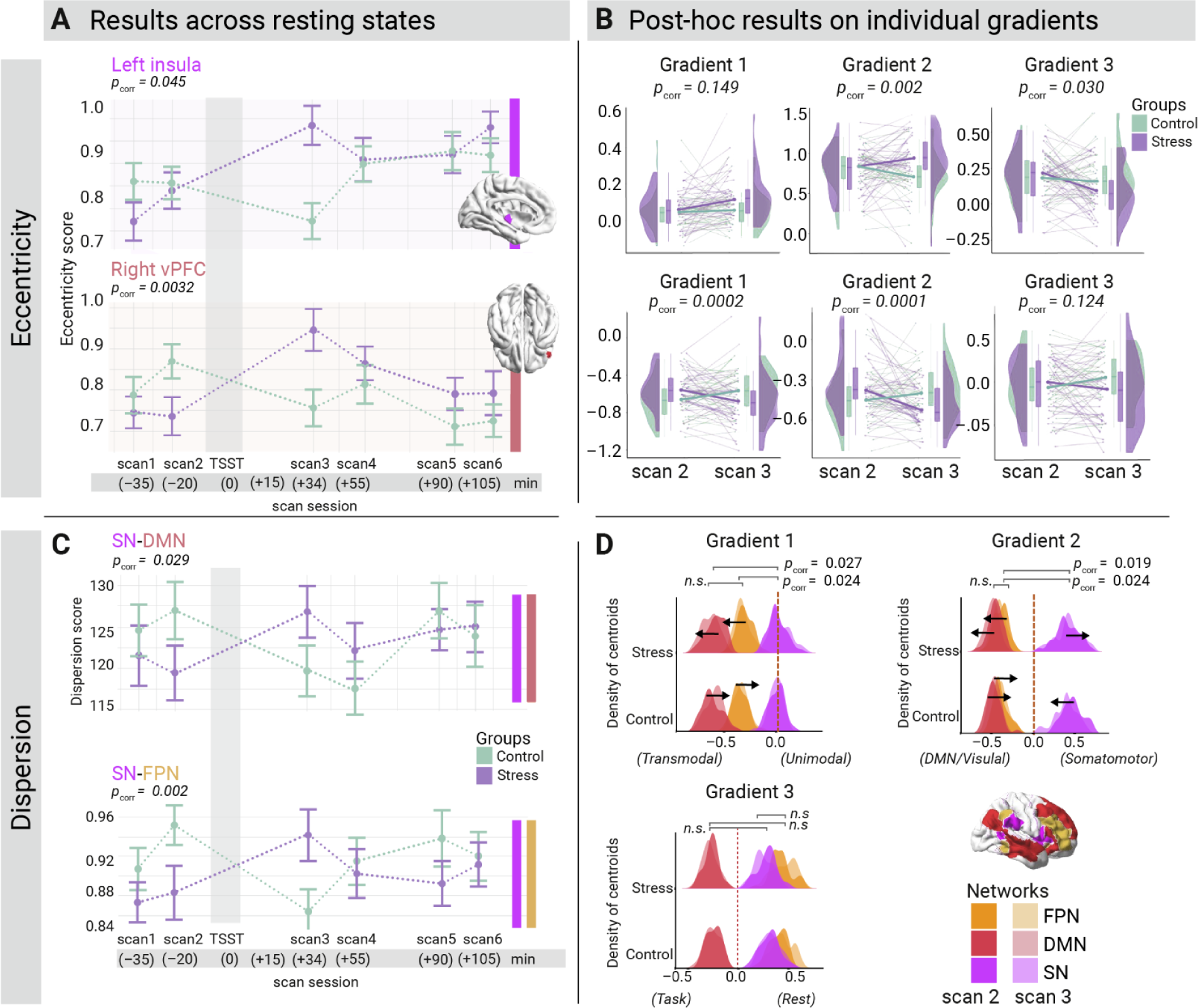
Significant changes in two parcels and in the network dispersion. (A) Eccentricity scores of right vPFC (DMN) and left insula (SN) across six resting states. We observed a significant interaction between rs-states 2 and 3 and stress compared to control groups. (B) Post hoc analysis: the left insula shifts to somatomotor processing, while the right vPFC shifts to DMN-like connectivity on the first two gradients. (C) Dispersion scores for significant network interactions across six resting states. A significant interaction between scans 2 and 3 and stress compared to the control group for the two network comparisons (SN-DMN) and (SN-FPN). (corr = Bonferroni corrected) (D) Density plot of centroids per gradient for the stress and control groups before and after stress. Along gradients 1 and 2, the FPN and DMN shifted significantly away from the SN (the distance between the centroids increased significantly) towards a more transmodal (DMN-like) connectivity profile. Along the second gradient, the SN shifted away from the other networks toward a somatomotor connectivity profile. (corr = FDR corrected).

Exploratory post-hoc analysis revealed significant interaction effects involving the left insula along gradient 2 (p_G2_ Bonferroni = 0.002) and gradient 3 (p_G3_ Bonferroni = 0.030) *(Figure 2B)* and a non-significant shift along gradient 1. Following the TSST, the stress group’s functional gradient connectivity profile showed a shift along gradient 2, becoming more similar to the somatomotor network. Simultaneously, along gradient 3, their connectivity profile shifted towards zero compared to their pre-TSST profile and the control group. In contrast, the control group exhibited an opposite shift along gradient 2 and maintained stability along gradient 3. The connectivity profile of the right vPFC of the stress group showed a shift towards the ‘transmodal’ end of gradient 1 (p_G1_ Bonferroni = 0.0002) and the ‘visual / DMN’ end of gradient 2 post-stressor (p_G2_ Bonferroni = 0.0001), while the control group showed an increase in gradient values along both gradients. We found no significant shift of the vPFC along gradient 3 (p_G3_ Bonferroni = 0.124).

### Dispersion

Within-network dispersion showed no significant interaction effects for the SN, FPN, and DMN (all p > 0.056, Bonferroni-corrected) in the context of acute stress. Between-network dispersion revealed significant interaction effects between the SN and DMN (p = 0.029, Bonferroni-corrected) and between the SN and FPN (p = 0.002, Bonferroni-corrected) *(Figure 2C)*. A post hoc analysis going beyond our preregistration revealed that in the stress group along gradient 1, the DMN and FPN decoupled from the SN, shifting towards the transmodal end of this gradient (p_G1 SN-DMN_ Bonferroni = 0.027; p_G1 SN-FPN_ Bonferroni = 0.024). Similarly, along gradient 2, the DMN and FPN decoupled from the SN, shifting towards the visual/DMN end, while the SN shifted towards the somatomotor end (p_G2 SN-DMN_ Bonferroni = 0.019; p_G2 SN-FPN_ Bonferroni = 0.024). We did not observe a significant association between the SN, FPN, and DMN along gradient 3 in our sample (all p Bonferroni > 0.96) *(Figure 2D)*.

### Peripheral and subjective stress measures: results

In the primary baseline-adjusted delta ANCOVAs, no association between changes in acute stress measures and changes in gradient-space eccentricity or dispersion was significant after multiple-comparison correction (Figure 3); all within-family FDR-adjusted p_FDR values were >= .962 for dispersion and >= .510 for eccentricity (Supplementary Figures 1 and 2). For dispersion, the smallest uncorrected p-value was observed for the DMN–SN distance, specifically for the Group × salivary cortisol change interaction p_uncorected_ = .097 (p_FDR_ = .962). For eccentricity, the smallest uncorrected p-value was observed for the association between vPFC eccentricity and salivary cortisol change, p_uncorected_ = .068 (p_FDR_ = .510), with HRV change, p_uncorected_ = .126 (p_FDR_ = .510). Exploratory Spearman rank correlations conducted within the stress group only also revealed no associations that survived within-family FDR correction; the strongest nominal effects were positive correlations of salivary cortisol change with insula eccentricity (ρ = .42, p_uncorrected = .018, p_FDR = .120) and vPFC eccentricity (ρ = .42, p_uncorrected = .020, p_FDR = .120), and a negative correlation between HRV change and SN–DMN dispersion (ρ = −.39, p_uncorrected = .037, p_FDR = .302)

**Figure 3.**
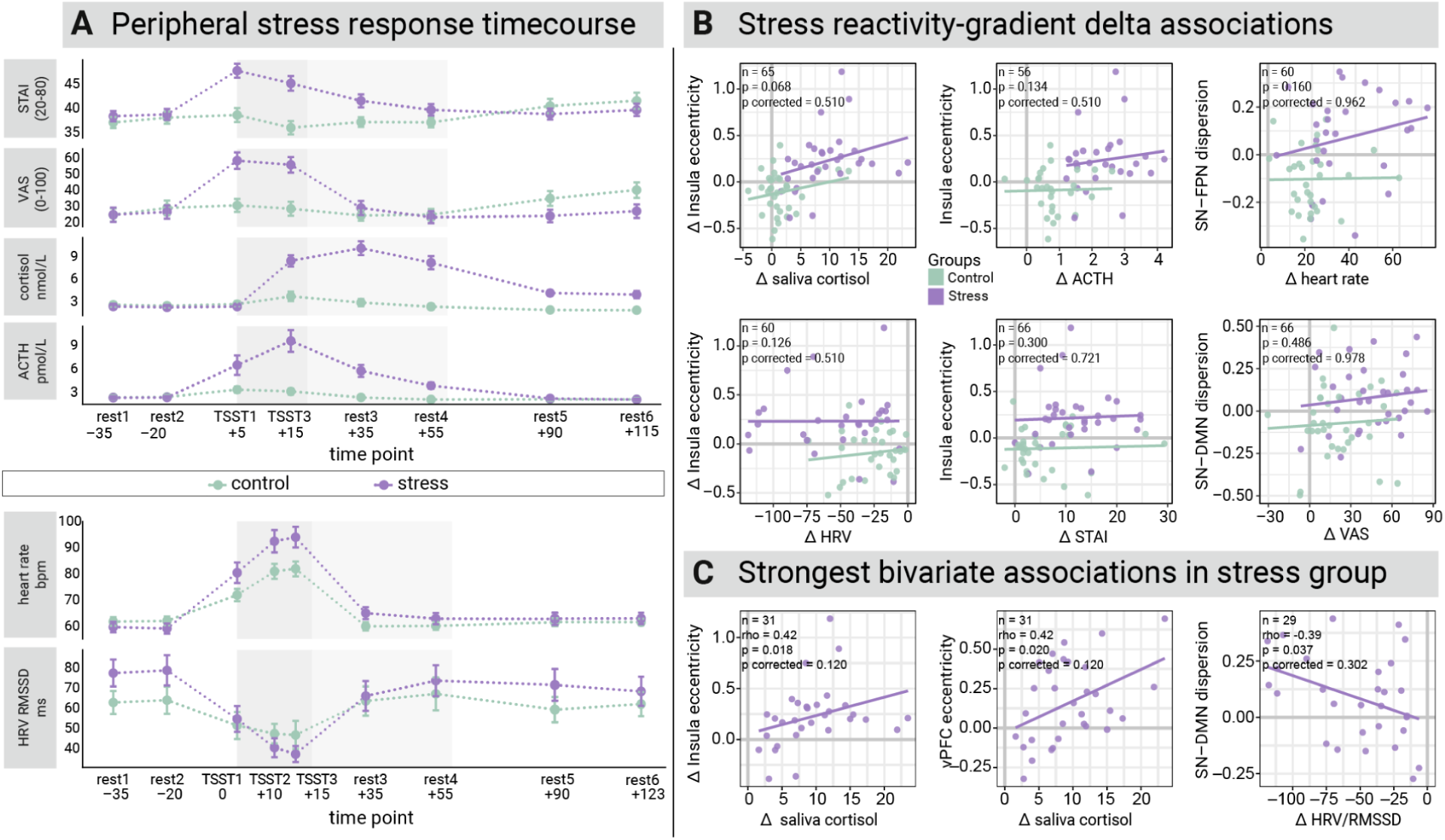
Peripheral stress responses across the experimental session and strongest exploratory associations with changes in brain functional organization. **(A)** (Top) Mean time courses of subjective and endocrine stress measures in the control (blue) and stress (orange) groups, including state anxiety (STAI; range 20–80), perceived stress (VAS; range 0–100), salivary cortisol (nmol/L), and plasma ACTH (pmol/L). To align these measures with the fMRI analysis epochs, questionnaire and endocrine data were averaged at the participant level into scan-relevant epochs using adjacent samples: rest1 = T3/T4, rest2 = T4/T5, TSST1 = T6, TSST3 = T7, rest3 = T8/T9, rest4 = T9/T10, rest5 = T11/T12, and rest6 = T12/T13 (displayed at −35, −20, +5, +15, +35, +55, +90, and +115 min relative to TSST onset, respectively); T14 was not included in the epoch-averaged display. The darker grey area marks the TSST period, and the lighter grey area marks the early post-stress interval during which participants in the stress group were still expecting a possible additional task. (Bottom) Mean time courses of autonomic stress measures, heart rate (bpm), and heart rate variability (HRV as root mean square of successive differences, RMSSD, in ms), across the same experimental session. Autonomic data were averaged within each recording segment (see Figure 1): rest1 = rest1a/rest1b, rest2 = rest2a/rest2b, TSST1 = anticipation, TSST2 = interview, TSST3 = arithmetic, rest3 = rest3a/rest3b, rest4 = rest4a/rest4b, and rest5 = rest5a/rest5b (displayed at −35, −20, 0, +10, +15, +35, +55, and +90 min). In A–B, points indicate group means, and error bars show standard errors of the mean. See (13–15) for more details. (B) Scatterplots of the strongest exploratory associations between peripheral stress reactivity and pre-to-post change in brain gradient measures, identified from baseline-adjusted delta ANCOVA models testing the Group × stress-measure interaction. For each peripheral measure (salivary cortisol, ACTH, heart rate, HRV, STAI, and VAS), the single brain outcome with the lowest uncorrected interaction *p* value is shown. Points represent individual participants, lines show group-specific fitted regression slopes, and panel annotations report the sample size and the uncorrected and corrected *p* values for the interaction term. (C) Scatterplots showing the strongest bivariate associations between peripheral stress responses and changes in brain gradient organization across participants. Greater increases in salivary cortisol were associated with larger increases in insula eccentricity and vPFC eccentricity, whereas larger decreases in HRV/RMSSD were associated with greater increases in SN–DMN dispersion. Points represent individual participants, and solid lines indicate fitted linear trends. Panel annotations report sample size, Spearman’s rho, and the corresponding uncorrected and multiple-comparison-corrected p values.

Supplementary three-way random-intercept LMM sensitivity analyses were also uniformly non-significant, with the strongest nominal effect for the DMN-SN distance with salivary cortisol, LRT p_uncorected_ = .121, providing no convergent support for robust stress-measure-to-gradient mapping effects in this dataset.

## Discussion

In the present study, we tested the preregistered (https://osf.io/5yspn) hypothesis of an association between acute psychosocial stress and functional connectivity profiles, in particular the organization of individual brain regions and large-scale cortical networks along functional cortical gradients. We also explored the relation of these brain changes with individual differences in subjective and peripheral stress reactivity. We used cortical gradient analysis, a data-driven method to map cortical functional hierarchies via low-dimensional summaries of resting-state functional connectivity (17). We first quantified the eccentricity of brain regions within gradient space, which reflects how functionally distinct each region is within the cortical hierarchy. We then quantified dispersion within and between the three stress-relevant large-scale cortical networks SN, FPN, and DMN, where higher within-network dispersion reflects greater functional heterogeneity among a network’s constituent regions, and higher between-network dispersion reflects greater functional segregation between networks within the cortical hierarchy (6,33). We found that acute stress was associated with a divergence in cortical connectivity profiles: the ventral prefrontal cortex, together with the DMN and FPN, shifted toward more transmodal connectivity, whereas the insula shifted toward more sensorimotor connectivity. In gradient terms, this means that ventral prefrontal, DMN, and FPN regions became functionally more similar to higher-order association cortex and less similar to unimodal sensory-motor cortex, while the insula showed the opposite pattern. Physiologically, this suggests stronger coupling among transmodal integrative regions on the one hand and tighter coupling of salience-and interoception-related insular processing with cortical systems involved in body-state representation and action readiness on the other. In our data, these changes in cortical functional organization were not significantly associated with measures of subjective or peripheral stress reactivity after correction for multiple comparisons. Yet, trend-level associations before correction pointed to potentially meaningful brain-body relationships: higher salivary cortisol was related to greater vPFC and insula eccentricity, and lower HRV was related to greater default mode–salience separation, showing that higher stress reactivity was linked to more pronounced adaptive cortical functional reorganization. Although these effects should be interpreted cautiously, they suggest that links between central and peripheral (autonomic, endocrine) stress dynamics may be present and warrant further investigation in future studies.

In the following, we discuss each finding in more detail:

### Shift of the right vPFC towards a DMN connectivity profile

Following acute stress, the right ventrolateral prefrontal cortex - a parcel spanning primarily orbital and ventrolateral regions but also portions of the inferior frontal gyrus (pars orbitalis) - shifted toward a DMN-like connectivity pattern along the first two gradients. In gradient space, the DMN occupies the extreme transmodal pole of cortical organization on all three cortical gradients, far from sensory and task-positive systems (17,18). A parcel shifting toward this anchor becomes more functionally similar to DMN nodes and more segregated from externally oriented networks (34,35). The ventrolateral PFC/inferior frontal gyrus (IFG) is traditionally linked to executive control and response inhibition (36)) and often assigned to the FPN/Control network (26); however, in the Schaefer parcellation, it falls within the DMN, consistent with accounts of an extended DMN (37–40). Meta-analyses show that acute stress robustly activates the vPFC/IFG and implicates it in affective appraisal and salience detection (41,42). Its post-stress shift toward the DMN may therefore reflect a decoupling from control networks and an alignment with DMN subsystems supporting self-referential thought and context integration (43) commonly found in post-stress reduction in cognitive control and heightened inward focus (44).

### Left insula shift toward somatomotor processing

Following acute stress, the left anterior insula parcel shifted along gradient 2 toward the somatomotor end of the cortical hierarchy. On gradient 2, this direction reflects increasing functional similarity to primary sensorimotor areas and decreasing functional similarity to transmodal regions (34,35). The shift likely reflects a stress-induced prioritization of bodily signal processing: the (anterior) insula, as a hub for interoceptive processing (45), may increase functional coupling with sensorimotor circuits to enhance motor readiness and sensorimotor integration. This is consistent with the anterior insula’s role in alerting attentional networks and gating conscious access to bodily states, facilitating swift, goal-directed actions essential for adapting to acute stress (46–48).

### Changes in tri-network interactions

Acute stress altered interactions among the three large-scale networks compared to before the stress intervention and compared to the control condition: The SN maintained its central position along the unimodal-transmodal axis (gradient 1), but shifted toward a somatomotor profile on gradient 2, while the DMN and FPN decoupled from the SN along both gradients, adopting more transmodal (DMN-like) connectivity. This spatial separation reflects increased between-network segregation in whole-cortex functional connectivity, extending prior reports of stress-related tri-network reorganization (6,49,50). For instance, prior findings of reduced SN coupling with the DMN and FPN under stress were interpreted as reflecting a functional divergence between networks: the SN amplifies salience detection and bodily readiness, whereas the DMN engages more strongly in self-reference (50). Notably, we observed no significant changes in within-network dispersion, indicating that, in the macroscale connectivity perspective, stress primarily reconfigures interactions *between* networks rather than *within* them. Overall, our findings suggest that acute stress leads to an adaptive segregation: the SN shifts toward sensorimotor circuits supporting rapid actions, while the FPN and DMN consolidate transmodal connectivity supporting self-referential processing and higher-order cognition, respectively.

### Linking central and peripheral dynamics

No association between gradient changes and peripheral stress reactivity survived multiple-comparison correction. The strongest nominal effects were a positive correlation between salivary cortisol and vPFC and insula eccentricity, and a negative association between HRV and SN-DMN dispersion. This pattern tentatively aligns with our previous findings (13–15), suggesting that peripheral stress measures may capture partly distinct facets of the central stress response, with salivary cortisol more closely linked to local (node-level) gradient changes and HRV to network-level reconfiguration.

## Limitations

Functional gradients compactly summarize macroscale connectivity, but, like other dimensionality-reduction approaches, they trade biological specificity for abstraction and are sensitive to analytic choices (51). Mechanistic interpretation should therefore be made carefully, and future studies would benefit from complementing gradient analysis with task-based activation or seed-based connectivity approaches. Further, gradient analysis is most established for the cerebral cortex (16,32) and cortical networks whose stress-related reconfiguration is well documented (6,49,50). Subcortical and cerebellar regions are comparatively less mature, though important advances now map gradients in the subcortex and cerebellum and their correspondence to cortical axes (52,53). Yet including subcortical organization under stress would be especially informative because core effectors of the stress response (e.g., hypothalamus, amygdala) are subcortical structures involved in orchestrating the network-level shifts observed after stress (54–56). Moreover, given their direct involvement in autonomic and endocrine regulation, stress-induced gradient changes in subcortical regions may be stronger candidates for associations with peripheral stress measures such as cortisol and HRV than cortical reorganization alone.

The sample was restricted to young healthy males, limiting generalizability to women and older adults who may differ systematically in autonomic, central, and endocrine stress reactivity. The moderate sample size further constrains statistical power for individual-differences analyses. For other specific limitations of the data set and collection procedure, see (13–15).

## Conclusion

Using cortical gradients, we evaluated how acute psychosocial stress reconfigures macroscale functional cortical organization. Acute stress elicited a selective, gradient-aligned reconfiguration, most pronounced in the right ventrolateral prefrontal cortex and left anterior insula, consistent with stress-driven shifts toward transmodal and somatomotor connectivity, respectively - likely reflecting heightened interoceptive processing, motor readiness, and self-referential reappraisal. Together, these findings situate acute stress effects within cortical gradient space. Trend-level associations between stress-related changes in cortical gradient space and salivary cortisol and HRV tentatively suggest that peripheral stress dynamics may be reflected in cortical gradient reorganization. This relationship remains to be confirmed in larger, more diverse samples. More broadly, these results demonstrate the utility of cortical gradient analysis as a sensitive framework for characterizing stress-induced reorganization of the brain’s functional hierarchy and motivate its integration with peripheral physiological measures in future stress neuroscience research.

## Supporting information

Supplementary_results

## Data and code availability

All derived data products—including cortical gradient values (eccentricity, dispersion) and peripheral physiological measures (HRV/RMSSD, cortisol, ACTH)—together with the analysis code, are openly available at https://doi.org/10.5281/zenodo.19920052 under a Creative Commons Attribution 4.0 International Licence (CC BY 4.0; https://creativecommons.org/licenses/by/4.0/).

## Author contributions

M.U., J.R., A.V., and M.G. conceived of the experiment. M.U., J.R., and M.G. collected the data. E.R. and A.P. wrote the analysis code and analyzed the data. A.V., M.U., S.H., and M.G. advised analyses. A.P. and E.R. wrote the manuscript, and all authors contributed to its editing. M.G. and A.V. secured funding for the project.

## Acknowledgements

Support: This work was supported by the German Federal Ministry of Education and Research (BMBF) under grants 13GW0206, 13GW0488, 16SV9156, the Deutsche Forschungsgemeinschaft (DFG) under grants 502864329 and 542559580, and by the cooperation between the Max Planck Society and the Fraunhofer-Gesellschaft (project NEUROHUM). (to ER and MG). During the work, AP was a pre-doctoral fellow of the International Max Planck Research School on the Life Course (LIFE, www.imprs-life.mpg.de; participating institutions: MPI for Human Development, Freie Universität Berlin, Humboldt-Universität zu Berlin, University of Michigan, University of Virginia, University of Zurich). During the work ER was a pre-doctoral fellow of the International Max Planck Research School on Cognitive Neuroimaging (CoNi, https://imprs-coni.mpg.de/)

Thanks: We thank Sofie Falk and Jonathan Smallwood for their advice.

