## Supplementary_results for "Central and peripheral dynamics of acute stress: evidence from functional cortical gradients"

### 1.1 Cross-atlas anatomical comparison of significant parcels

| Atlas | Atlas ID | Region | % of ROI | Category |
|---|---|---|---|---|
| **Brainnetome** | 163 | G_L | 39.1% | Hypergranular insula |
| | 169 | vld/vlg_L | 28.5% | Ventral dysgranular and granular insula |
| | 73 | TE1.0/TE1.2_L | 17.5% | Heschl’s gyrus |
| | 171 | dIg_L | 6.4% | Dorsal granular insula |
| | 77 | A38l_L | 2.0% | Lateral area 38 |
| | 157 | A1/2/3tonIa_L | 0.5% | Postcentral gyrus |
| | 79 | A22r_L | 0.3% | Superior temporal gyrus |
| **AAL** | 3001 | Insula_L | 45.1% | Insula |
| | 8101 | Temporal_Sup_L | 34.0% | Temporal / Auditory |
| | 8111 | Heschl_L | 6.1% | Auditory |
| | 2331 | Rolandic_Oper_L | 1.0% | Operculum |
| **Harvard-Oxford** | – | Insular Cortex | 71.7% | Insula |
| | – | Planum Polare | 25.5% | Temporal / Auditory |
| | – | Central Opercular Cortex | 1.45% | Operculum |
| | – | Heschl's Gyrus | 1.3% | Auditory |

**Supplementary Table 1: Cross-atlas anatomical composition of Schaefer’s 400 parcellation – ROI100 (7Networks_LH_SalVentAttn_FrOperIns_4).** The table summarizes overlap with the Brainnetome, AAL, and Harvard-Oxford atlases, showing Atlas ID, region name, percentage of ROI, and anatomical category.

| Atlas | Atlas ID | Region | % of ROI | Category |
|---|---|---|---|---|
| **Brainnetome** | 52 | A12/47l_R | 45.96% | Pars orbitalis/ Inferior frontal gyrus/ Ventrolateral PFC |
| | 36 | A45r_R | 25.13% | Pars triangularis/ Inferior frontal gyrus / Ventrolateral PFC |
| | 38 | A44op_R | 14.69% | Pars opercularis/Inferior frontal gyrus / Ventrolateral PFC |
| | 44 | A12/47o_R | 7.18% | Pars orbitalis/ Inferior frontal gyrus/ Ventrolateral PFC |
| | 168 | dIa_R | 1.15% | Dorsal insula |
| | 34 | A45c_R | 0.10% | Pars triangularis/ Inferior frontal gyrus / Ventrolateral PFC |
| | 166 | vIa_R | 0.08% | Ventral insula |
| | 32 | IFS_R | 0.05% | Inferior frontal sulcus |
| | 78 | A38l_R | 0.05% | Superior temporal gyrus / Insula |
| **AAL** | 2322 | Frontal_Inf_Orb_R | 73.14% | Ventrolateral PFC / Orbital gyrus |
| | 2312 | Frontal_Inf_Tri_R | 18.20% | Ventrolateral PFC / Inferior frontal gyrus |
| | 3002 | Insula_R | 8.49% | Insula |
| | 2212 | Frontal_Mid_Orb_R | 0.13% | Orbital / Middle frontal gyrus |
| **Harvard-Oxford** | – | Frontal Pole | 44.30% | Prefrontal cortex / frontal pole |
| | – | Frontal Orbital Cortex | 37.56% | Orbital frontal cortex |

| | | | | |
|---|---|---|---|---|
| | – | Inferior Frontal Gyrus, pars triangularis | 14.35% | Ventrolateral PFC / operculum |
| | – | Frontal Opercular Cortex | 3.78% | Operculum / inferior frontal cortex |

**Supplementary Table 2:** Cross-atlas anatomical composition of Schaefer's 400 parcellation – ROI 377 (7Networks_RH_Default_PFCv_3). The table summarizes overlap with the Brainnetome, AAL, and Harvard-Oxford atlases, showing Atlas ID, region name, percentage of ROI, and anatomical category.

### 1.2. Highest nominal associations between changes (delta) in brain eccentricity and each of the peripheral stress measures peak reactivity

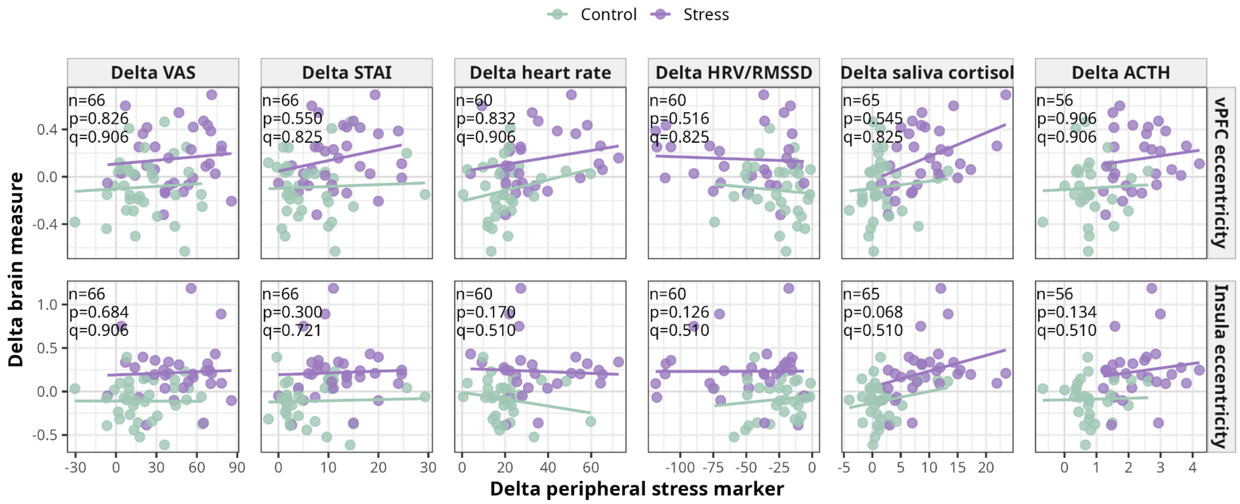


**Supplementary Figure 1.** Associations between changes in peripheral stress measures and changes in brain eccentricity. Panels show raw delta-versus-delta scatterplots for ventromedial prefrontal cortex (vPFC) eccentricity and insula eccentricity as a function of changes in VAS, STAI, heart rate, HRV/RMSSD, saliva cortisol, and ACTH. Blue points represent controls and orange points represent the stress group; lines indicate group-specific linear fits. In-panel text shows sample size (*n*), nominal *p* value, and FDR-corrected *q* value.

### 1.3. Highest nominal associations between changes (deltas) in the brain between network dispersion and each of the peripheral stress measures peak reactivity

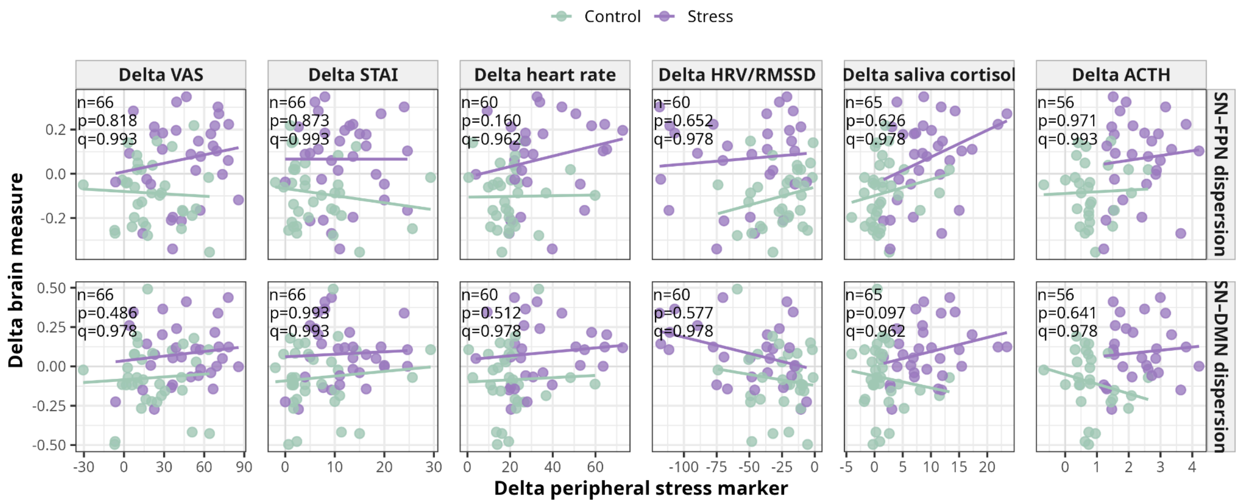


**Supplementary Figure 2:** Associations between changes in peripheral stress measures and changes in large-scale network dispersion. Panels show raw delta-versus-delta scatterplots for salience network–frontoparietal network (SN–FPN) dispersion and salience network–default mode network (SN–DMN) dispersion as a function of changes in VAS, STAI, heart rate, HRV/RMSSD, saliva cortisol, and ACTH. Blue points represent controls and orange points represent the stress group; lines indicate group-specific linear fits. In-panel text shows sample size (*n*), nominal *p* value, and FDR-corrected *q* value.
